# Genetic variation at the *Cyp6m2* putative insecticide resistance locus in *Anopheles gambiae* and *Anopheles coluzzii*

**DOI:** 10.1101/2020.11.12.378943

**Authors:** Martin G. Wagah, Petra Korlević, Christopher Clarkson, Alistair Miles, The *Anopheles gambiae* 1000 Genomes Consortium, Mara K. N. Lawniczak, Alex Makunin

## Abstract

**Background:** The emergence of insecticide resistance is a major threat to malaria control programmes in Africa, with many different factors contributing to insecticide resistance in its vectors, *Anopheles* mosquitoes. *CYP6M2* has previously been recognized as an important candidate in cytochrome P450-mediated detoxification in *Anopheles* mosquitoes. As it has been implicated in resistance against pyrethroids, organochlorines and carbamates, its broad metabolic activity makes it a potential agent in insecticide cross-resistance. Currently, allelic variation within the *Cyp6m2* gene remains unknown.

**Results:** Here, we use Illumina whole-genome sequence data from Phase 2 of the *Anopheles gambiae* 1000 Genomes Project (Ag1000G) to examine genetic variation in the *Cyp6m2* gene across 16 populations in 13 countries comprising *Anopheles gambiae* and *Anopheles coluzzii* mosquitoes. We find 15 missense biallelic substitutions at high frequency (defined as >5% frequency in one or more populations), that fall into five distinct haplotype groups that carry the main high frequency variants: A13T, D65A, E328Q, Y347F, I359V and A468S. We examine whether these alleles show evidence of selection either through potentially modified enzymatic function or by being linked to variants that change the transcriptional profile of the gene. Despite consistent reports of *Cyp6m2* upregulation and metabolic activity in insecticide resistant Anophelines, we find no evidence of directional selection occurring on these variants or on the haplotype clusters in which they are found.

**Conclusion:** Our results imply that emerging resistance associated with *Cyp6m2* is potentially driven by distant regulatory loci such as transcriptional factors rather than by its missense variants, or that other genes are playing a more significant role in conferring metabolic resistance.

## Background

Malaria remains a pernicious public health problem that plagues the African region, which has over 90% of the world’s malaria cases and deaths [1]. Although concerted vector control interventions such as long lasting insecticidal nets (LLINs) and indoor residual spraying (IRS) have led to the attainment of key milestones, global progress has stagnated and case numbers are stable or on the rise in many countries in Africa [1–3]. This is due to multiple factors, including the emergence of insecticide resistance, which threaten the effectiveness of vector control interventions [4].

The most well understood mechanisms of insecticide resistance are classified into two main functional categories depending on the underlying genes involved: target-site insensitivity and metabolic sequestration and detoxification. Both types may occur concurrently within a single population or even within a single mosquito [5–7]. These mechanisms have led to increasing resistance to all four common insecticide classes - pyrethroids, organochlorines, carbamates and organophosphates - in all major malaria vectors across Africa [7, 8].

Metabolic detoxification occurs mainly through the elevated activity of large and functionally diverse multigene enzyme families: glutathione S-transferases (GSTs), carboxylesterases (COEs) and cytochrome P450 monooxygenases (P450s) [7, 9]. Although a few candidates in these enzyme families have been directly associated with resistance, our understanding of metabolic resistance has lagged far behind that of pyrethroid target-site resistance, chiefly due to its complexity and the lack of associated causal mutations [10]. This is despite the fact that metabolic resistance is often considered a greater threat to mosquito control [9], especially since the only widely accepted occurrence of malaria vector control failure was attributed to the elevated expression of resistance-associated P450s in *An. funestus* [11–13]. A comprehensive understanding of metabolic resistance must therefore involve disambiguating the roles that individual enzymes play and the genetic backgrounds that underlie their significance in vector populations.

The CYP6M2 enzyme exhibits complex insecticide metabolism associated with multiple binding modes for insecticides [14]. Its gene is located within a cluster of 14 Cyp6 P450 genes on chromosome 3R of *An. gambiae* [15], and is among the 111 known P450 genes across the *An. gambiae* genome [16, 17]. In this genomic region, *Cyp6m2* is nested within a sub-cluster of P450s containing *Cyp6m3* and *Cyp6m4* which have also been associated with xenobiotic detoxification[18].

*Cyp6m2* is notably one of the few specific P450s that have shown a consistent association with metabolic resistance [5]. Metabolic resistance is mainly assessed through transcriptional profiling of genes involved in xenobiotic detoxification. Transcriptomic experiments such as quantitative PCR and microarray assays have established a link between *Cyp6m2* overexpression and the resistance phenotype in field populations of *An. gambiae*, *An. coluzzii*, *An. arabiensis* and *An. sinensis,* irrespective of the presence of knock-down resistance (*kdr*) mutations such as L995F or L995S in the voltage gated sodium channel (VGSC) [5, 19-21]. In DDT resistant *An. gambiae* in Ghana, *Cyp6m2* has been found to be overexpressed 3.2 to 5.2-fold in combination with the upregulation of additional P450s like *Cyp6z2* [18]. In DDT resistant *An. coluzzii* collected in Benin, *Cyp6m2* was also found to be overexpressed 1.2 to 4.6-fold in combination with *Gste2* from the *GST* gene family and in the presence of fixed *kdr* alleles in the *Vgsc* gene [22]. In Nigeria, the 2.4 to 2.7-fold upregulation of *Cyp6m2* was found to be associated with high levels of permethrin resistance [5] and *An. gambiae* that exhibited a strong resistance to bendiocarb in In Côte d’Ivoire also had an elevated (up to 8-fold) expression of the *Cyp6m2* gene [20]. In the same study, transgenic expression of *Cyp6m2* in *Drosophila melanogaster* was shown to produce resistance to both DDT and bendiocarb. *In vivo* functional analysis of multi-tissue overexpression induced by genetic modification has also shown *Cyp6m2* to be sufficient in conferring resistance to permethrin and deltamethrin [23]. However, this overexpression also increased the mosquitos’ susceptibility to the organophosphate malathion. Collectively, these studies indicate that *Cyp6m2* can confer metabolic resistance against insecticides in 3 of the 4 known classes: both type I and type II pyrethroids [14, 18, 23], organochlorines [24], and carbamates [20]. It therefore has a high potential for cross-resistance, which may make the problem of malaria vector control even more intractable by limiting the options available to malaria control programs for insecticide rotation or combination. The negative cross-resistance associated with malathion hereby points to potential mitigating strategies [23].

The frequent association of *Cyp6m2* with insecticide resistance described above warrants further investigation into whether there is evidence of copy number variation (CNV) or missense mutations at the locus. CNVs have been implicated in augmenting gene dosage leading to increased transcription of metabolic enzymes [25, 26]. A genome-wide CNV analysis conducted on the Ag1000G dataset and described in detail elsewhere [25] found CNVs to be significantly enriched in metabolic resistance-associated gene families and to be undergoing positive selection. These CNVs were identified across P450s (such as *Cyp9k1* and at both the *Cyp6z3-Cyp6z1* and the *Cyp6aa1-Cyp6p2* gene clusters) and GSTs (at the *Gstu4-Gste3* cluster). However, CNVs across the *Cyp6m2* locus were found to be rare, even in populations that are known to exhibit *Cyp6m2-*mediated resistance [25]. This indicates that CNVs alone are not sufficient to explain the widespread occurrence of the *Cyp6m2-* associated resistance phenotype: additional factors such as allelic variation might contribute to resistance associated with *Cyp6m2* activity.

Allelic variation can play an additional role in P450-mediated resistance by modifying either enzyme catalytic activity or gene expression levels [27]. Allelic variation has been shown to be key in inducing high metabolic efficiency of *Cyp6P9b* and in conferring metabolic resistance to *An. funestus* [28]. Allelic variants in metabolic genes have also been identified to reliably and reproducibly associate with resistance, such as in *Cyp4J5* and *Coeae1d* in *An. gambiae,* and can serve as diagnostic markers of phenotypic resistance [29]. However, there is still a paucity of information about allelic variation associated with metabolic resistance when compared to the well-characterized target-site mutations [29]. Mutations that may modulate metabolic resistance by either altering function or modifying expression in *Cyp6m2* are yet to be described.

Following the consistent association of *Cyp6m2* with insecticide resistance in many populations, we examine whole-genome Illumina sequence data from phase 2 of the *Anopheles gambiae* 1000 Genomes Project (Ag1000G) [30] which consists of 1,142 wild-caught mosquitoes sequenced to a mean depth above 14×, and report a comprehensive analysis of genetic variation within the *Cyp6m2* gene. We also examine the wider haplotypes around *Cyp6m2* spanning across the *Cyp6m* sub cluster and the larger *Cyp6* supercluster for signatures of selection.

## Results

### Cyp6m2 non-synonymous nucleotide variation

Short-read whole-genome sequence data from the Ag1000G phase 2 data resource [30] were used to investigate genetic variation at the *Cyp6m2* locus across 16 populations of *An. gambiae* and *An. coluzzii* (n = 1,142 total individuals) collected between 2000 and 2012 [*Table 1, Additional file 1*]. The single nucleotide polymorphisms (SNPs) we studied here were discovered and QC’d using methods described elsewhere [31]. We focused on SNPs that change the amino acid sequence of the *CYP6M2* enzyme as they have a potential functional role in *Cyp6m2*-associated insecticide resistance (n = 193) *[Additional file 2]*. As putative resistance variants under selection pressure from insecticides are expected to increase in frequency over time, we subsequently computed allele frequencies for every non-synonymous SNP in each population with reference to species and country of origin. We filtered the list to focus only on those variants that were at high frequency within populations or across populations (defined as >5% frequency in one or more populations). In total, this resulted in 15 non-synonymous variants that we further explored [*Table 1*].

**Table 1.**
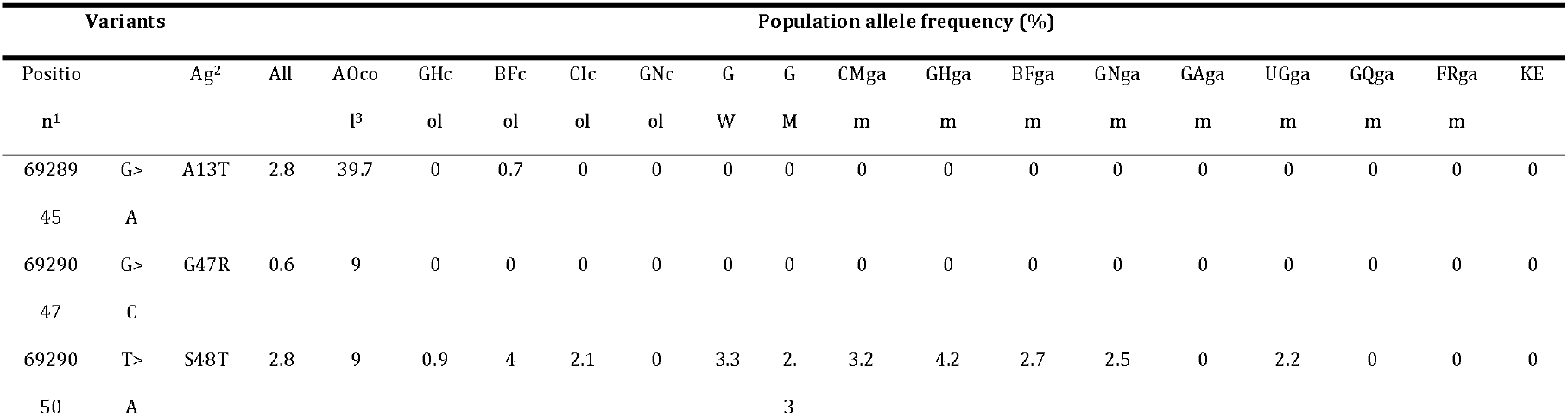
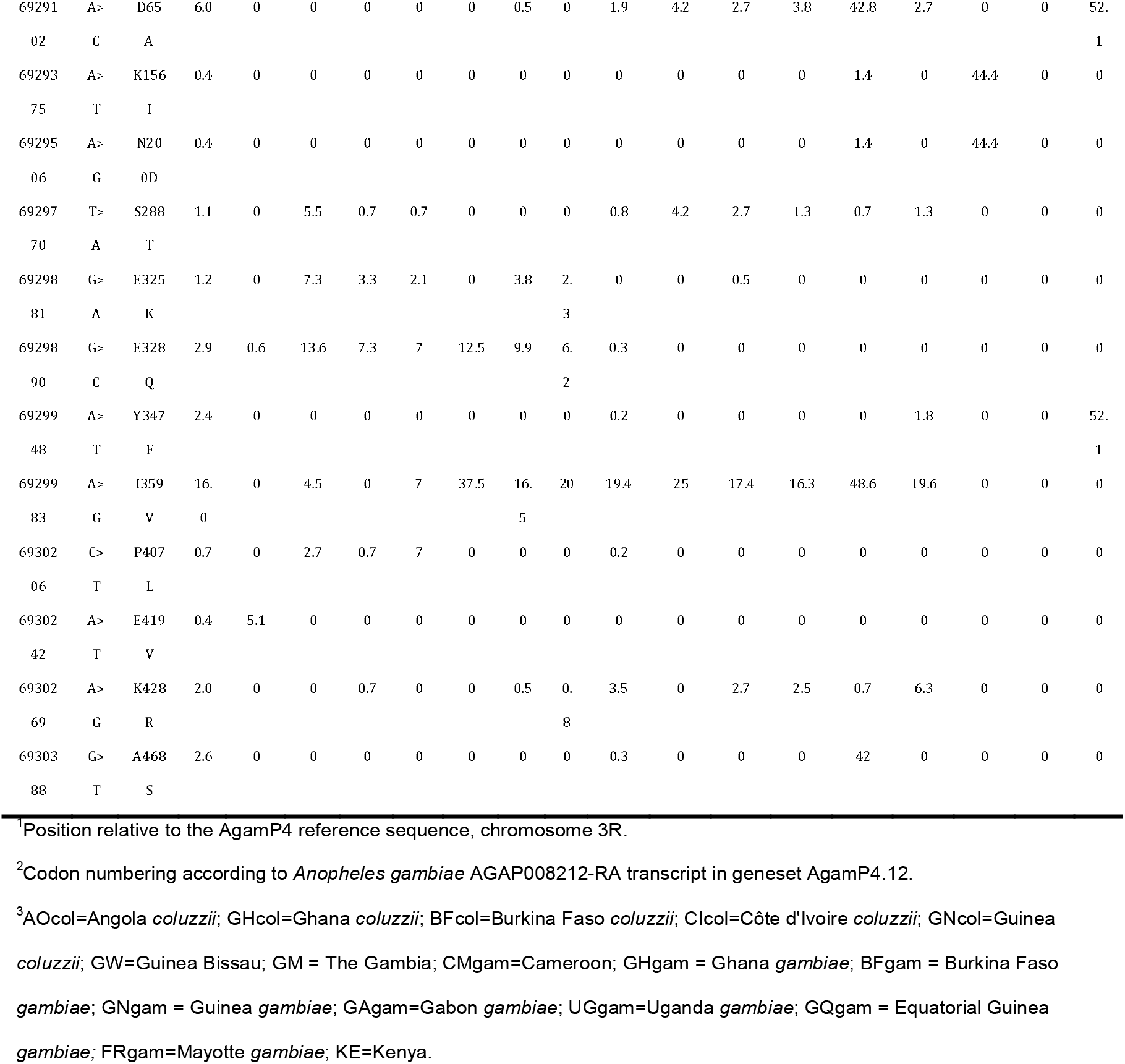
Allele frequencies of common *Cyp6m2* variants.

Analysis of the patterns of polymorphism of *Cyp6m2* from different populations showed both relative homogeneity within some geographical regions and distinct variants across different regions. The variants with the highest overall frequency were **I359V**(16%) and **D65A**(6%) [*Table 1*]. The most widespread variant was **I359V**, which was present in West, Central and East African populations of both *An. gambiae* and *An. coluzzii*. Populations with the highest frequency of **I359V**were Gabon (49%) and Ghana (25%) for *An. gambiae*, and Guinea (37.5%) for *An. coluzzii*. Another mutation, **E328Q**, was found across West Africa’s *An. coluzzii* populations in Burkina Faso, Côte d’Ivoire, Ghana, Guinea and The Gambia and ranged in frequency from 6.2 to 13.6%. Several variants were found to exceed the 5% threshold only in one or two populations: **A13T**and **Y347F,**in Angola’s *An. coluzzii* (39.7%) and in Kenya (52.1%) respectively and **D65A**only in Gabon’s *An. gambiae* and in Kenya’s populations at 42.8% and 52.1%, respectively [*Table 1*].

### Haplotypic backgrounds of non-synonymous alleles

The Ag1000G data resource contains data that not only spans across exonic regions of any given gene, but also intronic and intergenic regions. This enables a comprehensive analysis of haplotypes that contain putative insecticide resistance alleles, but is constrained by the fact that this resource does not contain samples whose resistance status or *Cyp6m2* expression levels are known.

Selection pressure acting upon missense variants or linked cis regulatory variants is likely to affect the haplotype structure of the gene. To study haplotype structure at *Cyp6m2*, we extracted biallelic SNPs across the entire 1689bp *Cyp6m2* gene to calculate the number of SNP differences between all pairs of 2,284 haplotypes derived from the mosquitoes. We identified a clustering threshold of seven SNPs where the haplotype clusters corresponded to the haplotypes carrying the high frequency alleles [*Table1*, *Figure 1*]. We found that these haplotypes could mostly be grouped into five distinct clusters (labelled C1-C5): C1 contained haplotypes carrying **A13T**; C2 contained most haplotypes carrying **D65A**, **A468S**, and some haplotypes carrying **I359V**; C3 contained most haplotypes carrying both **D65A** and **Y347F**, and C5 contained haplotypes carrying **E328Q. C4** contained haplotypes with no signature missense mutation [*Figure 1*].

**Figure 1.**
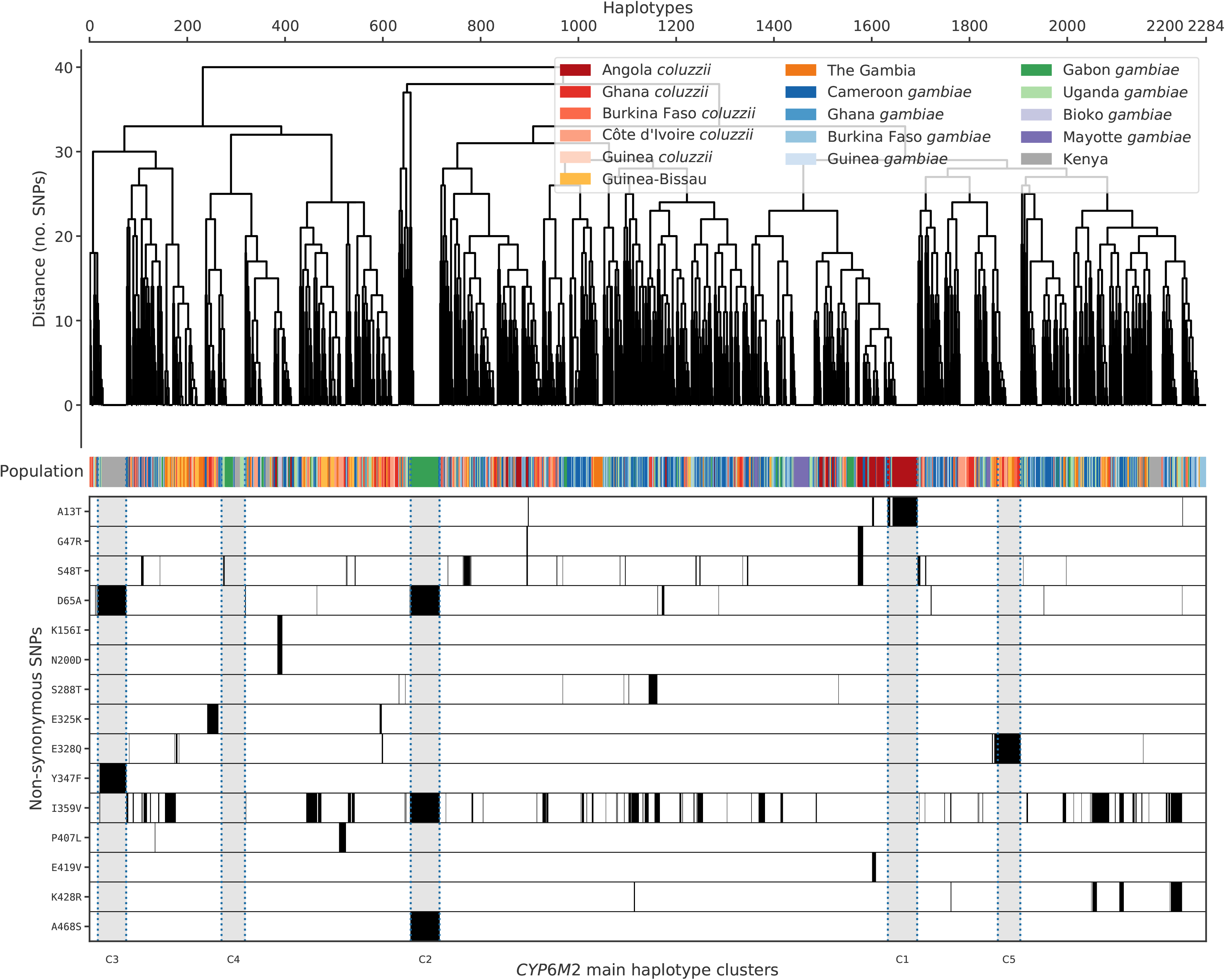
Hierarchical clustering of *Cyp6m2* haplotypes. Top: a dendrogram showing hierarchical clustering of haplotypes derived from wild-caught mosquitoes. The colour bar indicates the population of origin for each haplotype. Bottom: high frequency (>□5%) alleles identified within each haplotype (white = reference allele; black = alternative allele). The lowest margin labels the major haplotype clusters.

Overall, haplotype cluster distribution resembled the whole genome groupings of individuals described elsewhere using our dataset [30]: Cluster C5 contained haplotypes from West African *An. coluzzii*; C4 contained *An. gambiae* from West, Central and near-East Africa; and the rest of the clusters contained haplotypes from samples from a single country and species [*Figure 2*]. The variation across the haplotypes largely showed no strict or systematic difference between the two species or across broad geographic regions, which is in line with recent whole genome sequencing reports [31].

**Figure 2.**
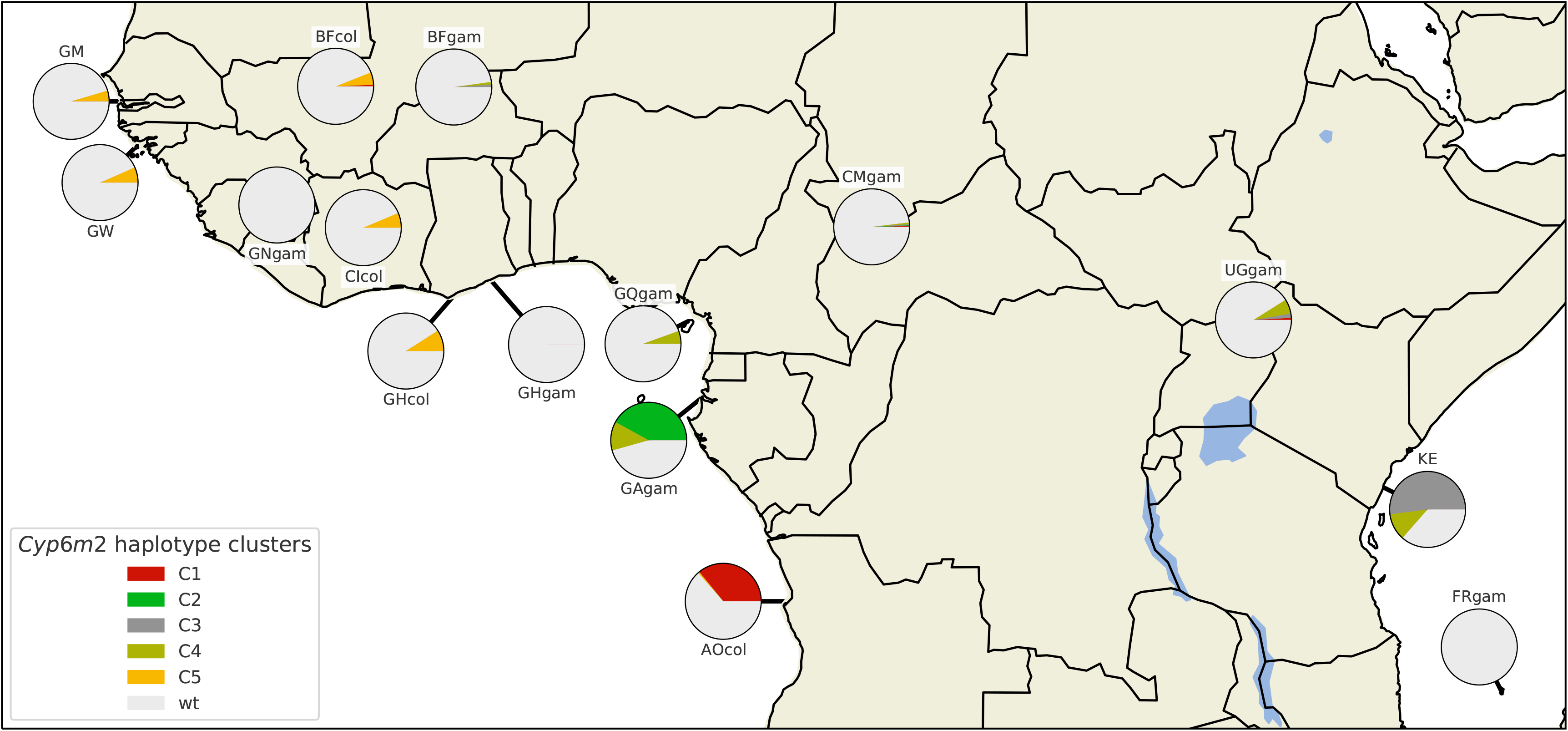
Map of haplotype cluster frequencies and distribution. Each pie chart indicates the haplotype group frequencies within specific sampling populations. The sizes of the wedges within the pies are proportional to haplotype group frequencies within the populations. Haplotypes in group C1 carry the **A13T** allele. Haplotypes in group C2 carry **D65A, I359V** and **A468S** alleles. Haplotypes in group C3 carry **D65A** and **Y347F** alleles. Haplotypes in group C5 carry the **E328Q** allele. Haplotypes in group C4 had no defining non-synonymous variant, and wild type (*wt*) haplotypes were all those that did not fall within the C1-C5 clusters.

We investigated patterns of association among these non-synonymous variants by computing the normalized coefficient of linkage disequilibrium (D’) using haplotypes from the Ag1000G phase 2 resource. Of the two highest frequency variants, **I359V** was found to be in perfect linkage with **A468S** but this was driven only by one population (Gabon) with most backgrounds carrying **I359V** not showing linkage with any other missense mutations [*Figure 1 & Supplementary Figure 1*]. **D65A** was in perfect linkage with **A468S** and **Y347F**, showing that **D65A** was almost only ever found on haplotypes carrying either **A468S** or **Y347F**. **I359V** and **D65A**, the highest frequency mutations across all populations, were found to be only in moderate linkage disequilibrium (0.36) [*Supplementary Figure 1*]. Other variants were found to be in weak linkage disequilibrium with the six main high frequency alleles and segregated independently within their own populations. While we observed some strong associations through linkage disequilibrium analysis across all populations, a deeper investigation revealed that these associations were driven by population specific dynamics in populations (such as Kenya) where we know bottlenecking has been an issue [31]. It is therefore unlikely that the identified variants are conferring some selective advantage against existing insecticide pressures.

We next explored whether the surrounding genomic region showed a similar hierarchical clustering pattern to *Cyp6m2*, which might be indicative of either dominant demographic effects or selection acting at other linked loci that is having a major impact on variation within *Cyp6m2*. The downstream genes we selected coded for proteins that were 1-to-1 orthologs with *D. melanogaster* genes. We selected *ODR2* [32], *HAM* [33] and *SH2* [34], which were 81280 bases, 457164 bases and 1198636 bases downstream of the *Cyp6m2* gene respectively. The distinctive haplotype clustering pattern observed for *Cyp6m2* in the Kenya, Angola and Gabon populations persisted across these genes, indicating that in these populations, the diversity reduction in and downstream of *Cyp6m2* is more likely driven by demography rather than by a selective sweep [*Supplementary Fig. 2-4*]. We also extracted biallelic SNPs across the *Cyp6m* sub cluster of 3 genes (*Cyp6m2*, *Cyp6m3* and *Cyp6m4*) and across the *Cyp6* supercluster of 14 genes within which the *Cyp6m* sub cluster is located (*Cyp6s2, Cyp6s1, Cyp6r1, Cyp6n2, Cyp6y2, Cyp6y1, Cyp6m1, Cyp6n1, Cyp6m2, Cyp6m3, Cyp6m4, Cyp6z3, Cyp6z2* and *Cyp6z1*), and performed hierarchical clustering across these regions as described above. The typical geographical stratification of haplotypes persisted, suggesting the absence of a selective sweep across this region [*Supplementary Fig.5 & 6*].

We examined the genetic backgrounds carrying these alleles further by constructing median joining networks (MJNs) [35] using the Ag1000G Phase 2 haplotype data. This enabled us to resolve the radiation of DNA substitutions arising on haplotypes carrying the identified variants. It also allowed us to reconstruct and position intermediate haplotypes while revealing the non-hierarchical relationships between haplotypes that could not be resolved by hierarchical clustering alone. The MJNs were constructed with reference to a maximum edge distance of two SNPs. This ensured that the connected components captured only closely related haplotypes. The resulting MJNs had a close correspondence with the hierarchical clustering output in assignment of haplotypes to clusters (88% overall concordance across all clusters).

The median joining networks showed more clearly the distinctive demographic stratification of the high frequency variants that was highlighted by the hierarchical clustering networks [*Figure 3*]. Most nodes containing secondary variants arising from the main nodes were small, which is inconsistent with directional selection where larger nodes are expected. Only one of the **I359V** nodes contained haplotypes from mosquitoes of both species, however the secondary nodes did not contain haplotypes from more than one species. This indicates that although **I359V** is shared by both *An. gambiae* and *An. coluzzii*, it is unlikely that this is because of an introgression event across the *Cyp6m2* gene.

**Figure 3.**
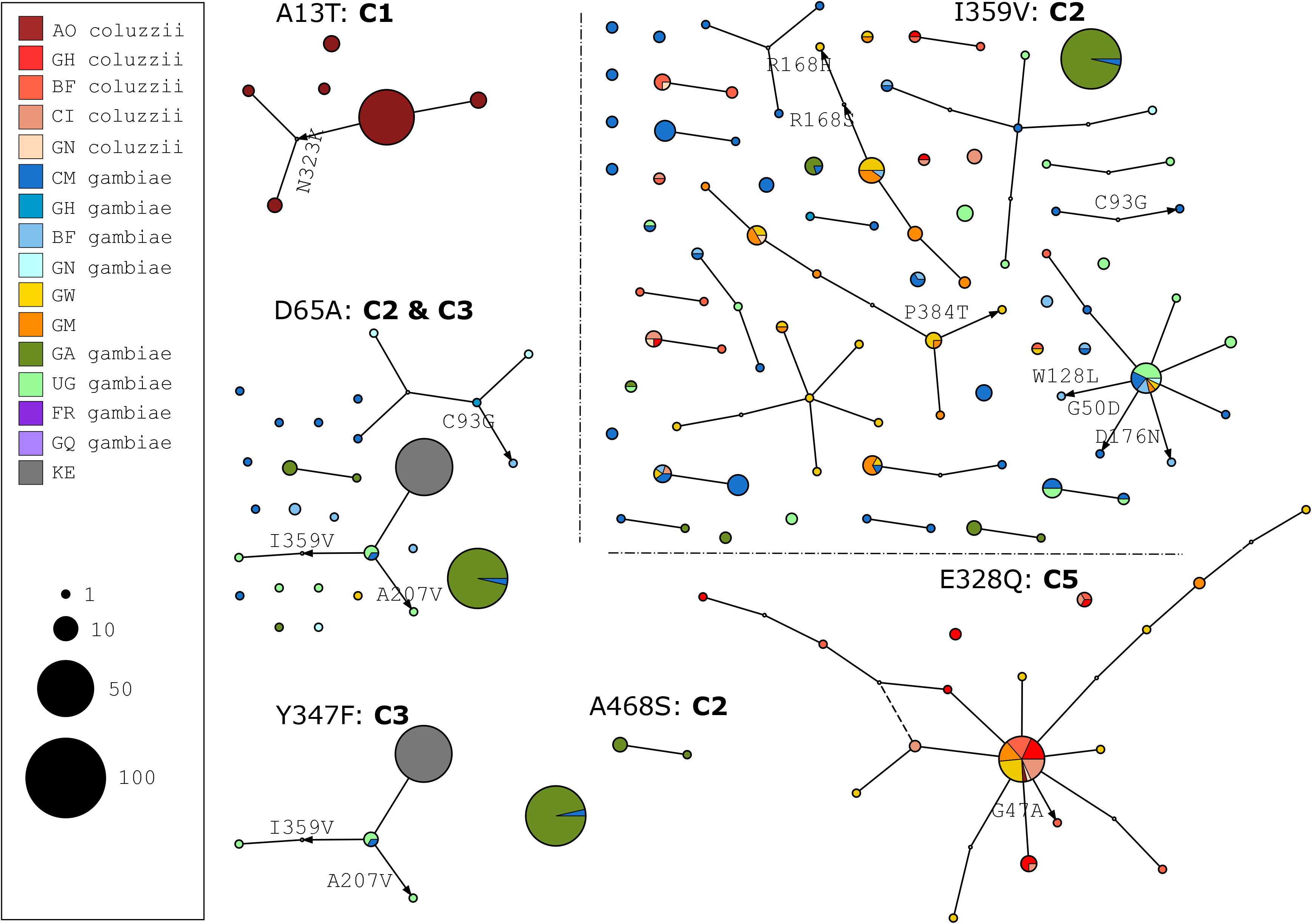
Haplotype networks. Median joining network for haplotypes carrying **A13T**, **D65A**, **E328Q**, **Y347F, I359V** and **A468S,** with a maximum edge distance of two SNPs. Node size indicates haplotype counts and node colour indicates the population/species of haplotypes. AO=Angola; GH=; BF=Burkina Faso; CI=Côte d’Ivoire; GN=Guinea; CM=Cameroon; GW=Guinea Bissau; GM = The Gambia; GA=Gabon; UG=Uganda; FR=Mayotte; GQ=Equatorial Guinea; KE=Kenya.

### Positive selection of non-synonymous alleles

Extended Haplotype Heterozygosity (EHH) decay [36] was calculated to explore evidence for directional selection on the haplotypes carrying high frequency non-synonymous variants. It is expected that the presence of ongoing or recent directional selection pressure would lead to the increase in frequency of haplotypes, which on average will have longer regions of haplotype homozygosity relative to haplotypes that are not under selection. This diversity reduction would produce signatures of selection that would be conspicuous across a large genomic region. EHH analysis would therefore be able to detect diversity reduction caused by ongoing directional selection being driven either by amino acid substitutions identified within the gene or by mutations within *cis*-acting elements next to the gene that may be under selection.

To perform the EHH decay analysis, we defined a core region of 1689 bases that spans across the entire gene. This was identical to what was used to differentiate the identified haplotype groups though hierarchical clustering. This region contained multiple distinct haplotypes above 1% frequency within the cohort, including haplotypes corresponding to the C1-C5 haplotype clusters. All haplotypes that did not correspond to C1-C5 were considered to be wild type (wt). Although there were several different haplotypes in each population that fit this description, we do not distinguish between them and call all these wild type, as *Cyp6m2* has no known resistance alleles and a true wild type remains to be discovered. EHH decay was then computed for each core haplotype up to 200 kilobases upstream and downstream [*Supplementary Fig. 7*]: beyond 200 kb, the EHH had decayed to zero.

We noted that haplotype clusters containing high frequency variants (C1-C5) did not exhibit a significantly slower EHH decay relative to the wild types, showing no evidence of positive selection. However, one Kenyan wild type haplotype group had a dramatically slower EHH decay relative to wild type haplotypes from other populations. In order to account for this difference within wild type groups across multiple populations and to reveal potential signs of selection that would be obscured by a collective analysis across all populations, we separated the haplotypes by population and species and recomputed EHH decay for each core haplotype as above.

Kenyan mosquito populations are known to have an extreme demographic history, as they have experienced a severe recent bottleneck, and the Angola and Gabon populations are known to be geographically unique populations which are strongly differentiated from all other populations[31]. Hence, their haplotypes exhibited a considerably slower decay than West African haplotypes [*first three panels: Figure 4*]. However, the putative resistance haplotypes C1-C5 did not experience a slower EHH decay relative to their wild type haplotypes, showing no evidence of positive selection acting upon those haplotypes in those populations.

**Figure 4.**
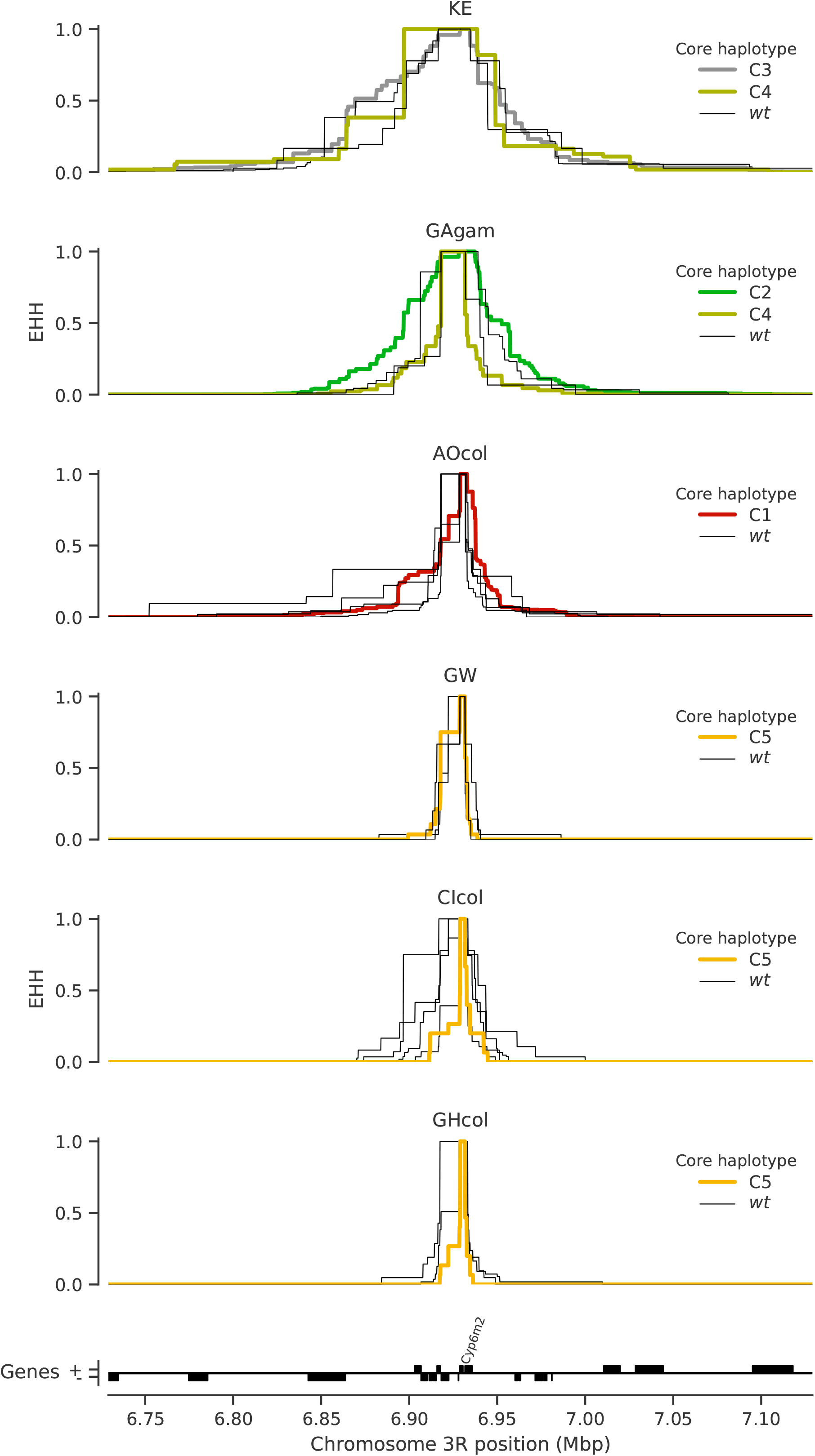
Extended haplotype homozygosity per population. No evidence for drastic difference in linkage disequilibrium within populations around core haplotypes across *Cyp6m2*. Extended Haplotype Heterozygosity (EHH) decay was calculated around cluster (C1 to C5) and non-cluster (wt) haplotypes using SNPs across and flanking the *Cyp6m2* region. KE=Kenya, GAgam=Gabon *An. gambiae*, AOcol= Angola *An. coluzzi*i, GW=Guinea Bissau, CIcol=Côte d’Ivoire *An. coluzzii*, GHcol=Ghana *An. coluzzii*.

As expected, the West African *An. coluzzii* haplotypes exhibited a much faster decay of EHH than specimens from Kenya, Angola, or Gabon, highlighting the demographic differences previously observed for these collections [31] [*last three panels, Figure 4*]. The C5 haplotype was a promising candidate for potential selection as it occurred within a more diverse population, and it was interesting to note that some wild type haplotypes in Côte d’Ivoire’s *An. coluzzii* had a slightly slower decay than others within West Africa [*fifth panel*, *Figure 4*]. However, these haplotypes were not part of the C5 cluster, and did not carry the widespread **E328Q** mutation. The C5 haplotype did not exhibit a dramatically slower decay of EHH than wild type haplotypes in the populations in which it was found, suggesting that it is not under positive selection.

## Discussion

*Cyp6m2* has been implicated in many *Anopheles* populations as a key P450 that contributes to the insecticide resistance phenotype [5, 14, 20, 24]. It has been reported that allelic variants across some P450s can affect enzyme conformational dynamics and substrate binding affinity [28], offering potential mechanisms that may modulate enzyme activity and efficiency, and thus account for additional *Cyp6m2* resistance where CNVs alone may not suffice. However, little is also known about *Cyp6m2* allelic variation across Africa.

In this study, we report a comprehensive account of the distribution of amino acid substitutions occurring within the *Cyp6m2* gene. We also examine the haplotype structure of the gene to probe for selective sweeps by performing hierarchical clustering of haplotypes. We also examine the genetic background upon which the missense variants are found by plotting both median joining networks and decay of extended haplotype homozygosity, which are useful for revealing signatures of selection. We note that the distinct haplotype groups therein are stratified demographically and largely correspond to signature missense variants found in specific populations. This is in contrast to the strong signals of recent positive selection at other cytochrome P450 gene loci such as at *Cyp6p3* [31] which is often upregulated in tandem with *Cyp6m2* in multiple pyrethroid resistant populations [5, 37, 38].

It is still unclear how the identified non-synonymous variants may modulate *Cyp6m2* binding activity, in either the presence or absence of multiple competitive substrates and metabolites. The two aromatic residues (Phe 108 and Phe 121) that have been previously identified to be vital in deltamethrin orientation in the *Cyp6m2* active site[14] were not found to contain high frequency variants in our dataset.

None of the haplotype groups identified that carried missense variants were found to be under directional selection. This is despite the existence of a widespread variant (**E328Q**) linked to a geographic region (West Africa) where *Cyp6m2* upregulation has been associated with emerging metabolic resistance [20, 37]. In *An. coluzzii* originating from both Côte d’Ivoire and Ghana, the C5 haplotype that carried **E328Q** was shown to have an even faster decay of EHH than the wild type haplotypes, further indicating an absence of directional selection. The stratification of other main haplotype clusters from Angola (C1), Gabon (C2) and Kenya (C3) was also consistent with the strong demographic differentiation and overall reduced heterozygosity of these populations described elsewhere [31].

While the genomic data quality across the *Cyp6m2* gene and its putative promoter region was satisfactory, there was a ~10,000 base region of inaccessibility upstream of *Cyp6m2* that cut across the intergenic region into *Cyp6n1* [39]. A similar inaccessible region was also present 1 kb downstream of the gene in the intergenic region between *Cyp6m2* and *Cyp6m3*, which is likely caused by the presence of repeats that inhibit read mapping. Although it is possible that the upstream region of inaccessibility could contain a regulatory variant that is susceptible to selection, it is unlikely to obscure signatures of selection.

It has been shown in multiple studies that target-site resistance (i.e. VGSC-*kdr*) provides a strong persistent baseline of resistance as it rises towards fixation within populations [40]. In the presence of insecticide selection pressure, target-site mutations and metabolic resistance have also been shown to act synergistically to confer a stronger resistance phenotype to pyrethroids [29, 41]. While signatures of selection have previously been identified in some metabolic gene clusters within populations that have a high *kdr* frequency[31], further studies need to examine whether directional selection occurring on one locus can obscure selection on another locus. To resolve this conundrum, genomic analysis must be performed on populations sampled across generations and whose transcriptomic and phenotypic characteristics are known, in order to tease out the individual contributions of specific sources of resistance.

Independent studies employing different experimental designs have also shown that metabolic resistance manifests as a cascade of multiple upregulated genes [42]. These genes, like *Cyp6m2*, are part of the normal cellular mechanism for xenobiotic detoxification that involves a linked, coordinated response of large multi-gene enzyme families in complicated pathways. Therefore, it is likely that identifying signatures of selection due to insecticide pressure will involve thorough analysis across this vast network. The Cap ‘n’ Collar isoform-C (CncC) transcription factor sub-family has been shown to work in tandem with other transcription factors to regulate the transcription of phase I, II and III detoxification loci of multiple insects such as *Culex quinquefasciatus* and *D. melanogaster* [43, 44]. *CncC* knockdown or upregulation has been shown to directly affect phenotypic resistance in *Anopheles gambiae* as well, modulating the expression of key P450s enzymes such as *Cyp6z2*, *Cyp6z3* and *Cyp6m2* that are located in the same genomic region[43, 44]. Given that we have detected no evidence of selection on amino acid variants in the *Cyp6m2* gene, it is possible that the emergence of *Cyp6m2* associated resistance is being driven by selection pressures acting upon genes coding for distant regulatory proteins such as transcription factors. These transcription factors can regulate downstream gene expression across large genomic distances. These transcription factors have also been implicated in the differential expression of other detoxification enzyme families also associated with insecticide resistance (GSTs, COEs, UDP-glucuronosyltransferases (UGTs) and ABC transporters). It is therefore likely that the centre of selection leading to the *Cyp6m2* associated resistance phenotype will be identified through whole genome selection scans of susceptible and resistant populations rather than by single loci analysis. Further research on Anopheline epigenomics, transcriptomics, proteomics and systems biology will also be game changers in mapping the complex regulatory network of insecticide resistance, aiding the identification of critical targets and the development of new strategies to control the spread of metabolic insecticide resistance.

## Conclusion

The scale up of insecticide-based interventions has caused increased selection pressure and higher levels of insecticide resistance across Africa. While the *CYP6M2* enzyme has been associated with emerging metabolic resistance in Africa, our data indicates that allelic variation within the *Cyp6m2* gene itself or across its Cyp6 supercluster has not been subject to recent positive selection in any of the populations sampled. This is in contrast to other Cytochrome P450 genes where CNV alleles are clearly under strong selection. Our results do not rule out a role for *Cyp6m2* in insecticide resistance in natural populations, but highlight the need for a deeper understanding of the regulatory networks affecting Cytochrome P450 gene expression in malaria vectors. This will require large-scale, holistic experimental work that collects genomic, transcriptomic and phenotypic datasets which when juxtaposed can resolve the complexities of metabolic resistance.

## Methods

### Data collection and analysis

In this study, we followed the species nomenclature of Coetzee *et al* [45] where *An. gambiae* refers to *An. gambiae sensu stricto* (S form) and *An. coluzzii* refers to *An. gambiae sensu stricto* (M form). A detailed description of the Ag1000G sample collection, DNA extraction, sequencing, variant calling, quality control and phasing can be found here [31]. Briefly, Anopheline samples were collected from 33 sampling sites across 16 populations in 13 countries in sub-Saharan Africa [*Table 1 & Additional file 1*]. The sampling procedure covered different ecosystems and aimed at collecting a minimum of 30 specimens per country. The specimens consisted of *An. gambiae* and *An. coluzzii*: only An. coluzzii were sampled from Angola, both *An. gambiae* and *An. coluzzii* were sampled from Burkina Faso, while all other populations consisted of *An gambiae,* except Kenya and Guinea Bissau where the species identity was indeterminate.

Whole genome sequencing of all mosquitoes was performed on the Illumina HiSeq 2000 platform. The generated 100 base paired-end reads were aligned to the *An. gambiae* AgamP3 reference genome assembly [46] and variants were called using GATK UnifiedGenotyper. Samples with mean coverage ⋜14× and variants with attributes that correlated with Mendelian error in genetic crosses were removed during quality control.

The SnpEff v4.1b software was used for the functional annotation of Ag1000G variant data [47] using locations from geneset AgamP4.12. All variants in transcript AGAP008212-RA with a SnpEff annotation of “missense” were regarded as nonsynonymous variants. The *Cyp6m2* gene has not been shown to exhibit alternative splicing, and no alternative transcripts have been reported.

### Haplotype clustering, linkage disequilibrium and mapping of haplotype clusters

To reveal the haplotype structure at *Cyp6m2, Cyp6m* sub-cluster, *Cyp6* supercluster, *HAM, ODR-2* and *SH2*, we computed the Hamming distance between all haplotype pairs and performed hierarchical clustering of haplotypes. We worked through arbitrary clustering threshold values to cut the dendrograms at genetic distances that would best highlight the most relevant clusters. We used Lewontin’s *D*′ [48] to compute the linkage disequilibrium (LD) between all pairs of missense *Cyp6m2* mutations. Image rendering for the haplotype clustering, linkage disequilibrium and haplotype cluster frequencies map was performed using the matplotlib Python package [49]. Geography handling for the haplotype cluster frequencies map was done using cartopy [50].

### Haplotype Networks

We constructed haplotype networks using the median-joining algorithm [35] implemented in Python [51]. Haplotypes carrying the main high frequency mutations were analysed with a maximum edge distance of two SNPs. The Graphviz library was used to render the networks and the composite figure was constructed in Inkscape [52].

### Extended haplotype homozygosity

We defined the core haplotype on a 1689 base region spanning the *Cyp6m2*, from chromosome arm 3R, starting at position 6928858 and ending at position 6930547. We selected this region to ensure a 1:1 haplotype correspondence with that used in the hierarchical clustering analysis. We computed extended haplotype homozygosity (EHH) across all core haplotypes in all populations as described in Sabeti et al. [36] using scikit-allel version 1.1.9 [53]. EHH composite plots were made using the matplotlib Python package [49].

## Supporting information

Supplementary Figure 2. Hierarchical clustering and missense mutations for ODR2.

Supplementary Figure 3. Hierarchical clustering and missense mutations for HAM

Supplementary Figure 4. Hierarchical clustering and missense mutations for SH2.

Supplementary Figure 5. Hierarchical clustering and missense mutations for Cyp6m sub cluster.

Supplementary Figure 6. Hierarchical clustering and missense mutations for Cyp6 supercluster.

Supplementary Figure 7. Extended haplotype homozygosity across all populations.

## List of abbreviations

CncC: Cap ‘n’ Collar isoform-C
CNV: Copy Number Variation
COEs: Carboxylesterases
DDT: Dichlorodiphenyltrichloroethane
EHH: Extended Haplotype Heterozygosity
GSTs: Glutathione S-Transferases
HAM: Transcription Factor Hamlet
IRS: Indoor Residual Spraying
*kdr*: Knock-Down Resistance
LLINs: Long Lasting Insecticidal Nets
MJNs: Median-Joining Networks
ODR2: Odd-Skipped Related
SH2: SRC Homology 2
SNPs: Single Nucleotide Polymorphisms
P450s: Cytochrome P450 Monooxygenases
UGTs: UDP-glucuronosyltransferases
VGSC: Voltage Gated Sodium Channel
*wt*: wild type

## Declarations

## Acknowledgements

The authors would like to thank the staff of the Wellcome Sanger Institute Sequencing and Informatics facilities for their contributions.

## Availability of data and materials

Jupyter Notebooks and scripts containing all analyses, tables and figures can be found in the GitHub repository [51]. Variant calls and phased haplotype data from the Ag1000G Phase 2 AR3 data release were used, and can be found here [54].

## Authors contribution

AM and MKNL designed the study. AM and CC developed the base code. MGW and AM performed all analyses. MGW drafted the manuscript. All authors read and approved the final manuscript.

## Consent for publication

Not applicable

## Ethics approval and consent to participate

Not applicable.

## Supplementary Figures

**Supplementary Figure 1. Linkage disequilibrium (*D*′) between non-synonymous variants.**

A value of 1 shows perfect linkage between the alleles. A value of −1 shows that the alleles are never found conjointly. The bar plot indicates allele frequencies within the Ag1000G phase 2 cohort.

**Supplementary Figure 2. Hierarchical clustering and missense mutations for *ODR2***.

Top: a dendrogram showing hierarchical clustering of haplotypes across the ODR2 gene. The gene is located at position 7,059,422 to 7,119,244: 128,875 bases downstream of *Cyp6m2*.

The colour bar indicates the population of origin for each haplotype.

Bottom: high frequency (>□5%) alleles identified within each haplotype (white = reference allele; black = alternative allele).

**Supplementary Figure 3. Hierarchical clustering and missense mutations for *HAM***.

Top: a dendrogram showing hierarchical clustering of haplotypes across the HAM gene. The gene is located at position 7,435,306 to 7,485,012: 504,759 bases downstream of *Cyp6m2*.

The colour bar indicates the population of origin for each haplotype.

Bottom: high frequency (>□5%) alleles identified within each haplotype (white = reference allele; black = alternative allele).

**Supplementary Figure 4. Hierarchical clustering and missense mutations for *SH2***.

Top: a dendrogram showing hierarchical clustering of haplotypes across the SH2 gene. The gene is located at position 8,176,778 to 8,183,084: 1,246,231 bases downstream of *Cyp6m2*.

The colour bar indicates the population of origin for each haplotype.

Bottom: high frequency (>□5%) alleles identified within each haplotype (white = reference allele; black = alternative allele).

**Supplementary Figure 5. Hierarchical clustering and missense mutations for *Cyp6m* sub cluster**.

Top: a dendrogram showing hierarchical clustering of haplotypes across the Cyp6m sub cluster of genes containing Cyp6m2, Cyp6m3 and Cyp6m4. The genes are located at position 6928858 to 6935721.

The colour bar indicates the population of origin for each haplotype.

Bottom: high frequency (>□5) alleles identified within each haplotype (white = reference allele; black = alternative allele).

**Supplementary Figure 6. Hierarchical clustering and missense mutations for *Cyp6* supercluster**.

Top: a dendrogram showing hierarchical clustering of haplotypes across the Cyp6 supercluster of 14 P450 genes containing *Cyp6s2, Cyp6s1, Cyp6r1, Cyp6n2, Cyp6y2, Cyp6y1, Cyp6m1, Cyp6n1, Cyp6m2, Cyp6m3, Cyp6m4, Cyp6z3, Cyp6z2* and *Cyp6z1*. The genes are located at position 6903106 to 6978142.

The colour bar indicates the population of origin for each haplotype.

Bottom: high frequency (>□70%) alleles identified within each haplotype (white = reference allele; black = alternative allele).

**Supplementary Figure 7. Extended haplotype homozygosity across all populations**.

A rapid decay of EHH in comparison to other haplotypes implies absence of positive selection.

## Additional files

1. Additional file 1.

a. File name = Additional file 1
b. Title = List of *An. gambiae* and *An. coluzzii* genome samples and haplotypes from Ag1000G Phase 2-AR3.
c. Format = csv
d. Description = Table showing Ag1000G Phase 2-AR3 sample properties such as population, country, region, sex, species identity and haplotype cluster.
2. Additional file 2.

a. File name = Additional file 2
b. Title = List of synonymous and non-synonymous genetic variants in *Cyp6m2*.
c. Format = csv
d. Description = Table showing Ag1000G Phase 2-AR3 *Cyp6m2* variant calls and variant properties stratified by population and effect.

## References

1. WHO: World Malaria Report 2019. 2019.

2. Bhatt S, Weiss DJ, Cameron E, Bisanzio D, Mappin B, Dalrymple U, Battle K, Moyes CL, Henry A, Eckhoff PA et al: The effect of malaria control on Plasmodium falciparum in Africa between 2000 and 2015. Nature 2015, 526 (7572)207–211.

3. Cibulskis RE, Alonso P, Aponte J, Aregawi M, Barrette A, Bergeron L, Fergus CA, Knox T, Lynch M, Patouillard E et al: Malaria: Global progress 2000 - 2015 and future challenges. Infect Dis Poverty 2016, 5 (1):61.

4. Ranson H, Lissenden N: Insecticide Resistance in African Anopheles Mosquitoes: A Worsening Situation that Needs Urgent Action to Maintain Malaria Control. Trends Parasitol 2016, 32 (3)187–196.

5. Djouaka RF, Bakare AA, Coulibaly ON, Akogbeto MC, Ranson H, Hemingway J, Strode C: Expression of the cytochrome P450s, CYP6P3 and CYP6M2 are significantly elevated in multiple pyrethroid resistant populations of Anopheles gambiae s.s. from Southern Benin and Nigeria. BMC Genomics 2008, 9:538.

6. Djouaka R, Riveron JM, Yessoufou A, Tchigossou G, Akoton R, Irving H, Djegbe I, Moutairou K, Adeoti R, Tamò M et al: Multiple insecticide resistance in an infected population of the malaria vector Anopheles funestus in Benin. Parasit Vectors 2016, 9:453.

7. WHO: Global report on insecticide resistance in malaria vectors: 2010--2016. 2018.

8. WHO Malaria Threats Map [https://apps.who.int/malaria/maps/threats/?theme=prevention&mapType=prevention%3A0&bounds=%5B%5B-54.61667525407141%2C-26.993804332606665%5D%2C%5B66.07511128112793%2C35.549094294064915%5D%5D&insecticideClass=PYRETHROIDS&insecticideTypes=&assayTypes=MOLECULAR_ASSAY%2CBIOCHEMICAL_ASSAY%2CSYNERGIST-INSECTICIDE_BIOASSAY&synergistTypes=&species=&vectorSpecies=&surveyTypes=&deletionType=HRP2_PROPORTION_DELETION&plasmodiumSpecies=P._FALCIPARUM&drug=DRUG_AL&mmType=1&endemicity=false&countryMode=false&storyMode=false&storyModeStep=0&filterOpen=false&filtersMode=filters&years=2010%2C2018]

9. Ranson H, N’Guessan R, Lines J, Moiroux N, Nkuni Z, Corbel V: Pyrethroid resistance in African anopheline mosquitoes: what are the implications for malaria control? Trends Parasitol 2011, 27 (2)91–98.

10. Wilding CS, Weetman D, Steen K, Donnelly MJ: High, clustered, nucleotide diversity in the genome of Anopheles gambiae revealed through pooled-template sequencing: implications for high-throughput genotyping protocols. BMC Genomics 2009, 10:320.

11. Hargreaves K, Koekemoer LL, Brooke BD, Hunt RH, Mthembu J, Coetzee M: Anopheles funestus resistant to pyrethroid insecticides in South Africa. Med Vet Entomol 2000, 14 (2)181–189.

12. Wondji CS, Morgan J, Coetzee M, Hunt RH, Steen K, Black WCt, Hemingway J, Ranson H: Mapping a quantitative trait locus (QTL) conferring pyrethroid resistance in the African malaria vector Anopheles funestus. BMC Genomics 2007, 8:34.

13. Wondji CS, Irving H, Morgan J, Lobo NF, Collins FH, Hunt RH, Coetzee M, Hemingway J, Ranson H: Two duplicated P450 genes are associated with pyrethroid resistance in Anopheles funestus, a major malaria vector. Genome Res 2009, 19 (3)452–459.

14. Stevenson BJ, Bibby J, Pignatelli P, Muangnoicharoen S, O’Neill PM, Lian L-Y, Müller P, Nikou D, Steven A, Hemingway J et al: Cytochrome P450 6M2 from the malaria vector Anopheles gambiae metabolizes pyrethroids: Sequential metabolism of deltamethrin revealed. Insect Biochem Mol Biol 2011, 41 (7):492–502.

15. Chromosome 3R: 6,928,825-6,930,580 - Region in detail - Anopheles gambiae - VectorBase [https://www.vectorbase.org/Anopheles_gambiae/Location/View?db=core;g=AGAP008212;r=3R6928825-6930580;t=AGAP008212-RA]

16. Ranson H, Claudianos C, Ortelli F, Abgrall C, Hemingway J, Sharakhova MV, Unger MF, Collins FH, Feyereisen R: Evolution of supergene families associated with insecticide resistance. Science 2002, 298 (5591)179–181.

17. Ranson H, Paton MG, Jensen B, McCarroll L, Vaughan A, Hogan JR, Hemingway J, Collins FH: Genetic mapping of genes conferring permethrin resistance in the malaria vector, Anopheles gambiae. Insect Mol Biol 2004, 13 (4)379–386.

18. Müller P, Donnelly MJ, Ranson H: Transcription profiling of a recently colonised pyrethroid resistant Anopheles gambiae strain from Ghana. BMC Genomics 2007, 8:36.

19. Nardini L, Christian RN, Coetzer N, Ranson H, Coetzee M, Koekemoer LL: Detoxification enzymes associated with insecticide resistance in laboratory strains of Anopheles arabiensis of different geographic origin. Parasit Vectors 2012, 5:113.

20. Edi CV, Djogbénou L, Jenkins AM, Regna K, Muskavitch MAT, Poupardin R, Jones CM, Essandoh J, Kétoh GK, Paine MJI et al: CYP6 P450 enzymes and ACE-1 duplication produce extreme and multiple insecticide resistance in the malaria mosquito Anopheles gambiae. PLoS Genet 2014, 10 (3):e1004236.

21. Yan Z-W, He Z-B, Yan Z-T, Si F-L, Zhou Y, Chen B: Genome-wide and expression-profiling analyses suggest the main cytochrome P450 genes related to pyrethroid resistance in the malaria vector, Anopheles sinensis (Diptera Culicidae). Pest Manag Sci 2018, 74 (8)1810–1820.

22. Djègbè I, Agossa FR, Jones CM, Poupardin R, Cornelie S, Akogbéto M, Ranson H, Corbel V: Molecular characterization of DDT resistance in Anopheles gambiae from Benin. Parasit Vectors 2014, 7:409.

23. Adolfi A, Poulton B, Anthousi A, Macilwee S, Ranson H, Lycett GJ: Functional genetic validation of key genes conferring insecticide resistance in the major African malaria vector, Anopheles gambiae. Proc Natl Acad Sci U S A 2019, 116 (51)25764–25772.

24. Mitchell SN, Stevenson BJ, Müller P, Wilding CS, Egyir-Yawson A, Field SG, Hemingway J, Paine MJI, Ranson H, Donnelly MJ: Identification and validation of a gene causing cross-resistance between insecticide classes in Anopheles gambiae from Ghana. Proc Natl Acad Sci U S A 2012, 109 (16)6147–6152.

25. Lucas ER, Miles A, Harding NJ, Clarkson CS, Lawniczak MKN, Kwiatkowski DP, Weetman D, Donnelly MJ, Anopheles gambiae Genomes C: Whole-genome sequencing reveals high complexity of copy number variation at insecticide resistance loci in malaria mosquitoes. Genome Res 2019, 29 (8)1250–1261.

26. Weetman D, Djogbenou LS, Lucas E: Copy number variation (CNV) and insecticide resistance in mosquitoes: evolving knowledge or an evolving problem? Curr Opin Insect Sci 2018, 27 82–88.

27. Schuler MA, Berenbaum MR: Structure and function of cytochrome P450S in insect adaptation to natural and synthetic toxins: insights gained from molecular modeling. J Chem Ecol 2013, 39 (9)1232–1245.

28. Ibrahim SS, Riveron JM, Bibby J, Irving H, Yunta C, Paine MJI, Wondji CS: Allelic Variation of Cytochrome P450s Drives Resistance to Bednet Insecticides in a Major Malaria Vector. PLoS Genet 2015, 11 (10):e1005618.

29. Weetman D, Wilding CS, Neafsey DE, Müller P, Ochomo E, Isaacs AT, Steen K, Rippon EJ, Morgan JC, Mawejje HD et al: Candidate-gene based GWAS identifies reproducible DNA markers for metabolic pyrethroid resistance from standing genetic variation in East African Anopheles gambiae. Sci Rep 2018, 8 (1):2920.

30. Clarkson CS, Miles A, Harding NJ, Lucas ER, Battey CJ, Amaya-Romero JE, Cano J, Diabate A, Constant E, Nwakanma DC et al: Genome variation and population structure among 1,142 mosquitoes of the African malaria vector species *Anopheles gambiae* and *Anopheles coluzzii*. bioRxiv 2019:864314.

31. Consortium TAgG: Genetic diversity of the African malaria vector Anopheles gambiae. Nature 2017, 552 (7683)96–100.

32. Chromosome 3R: 7,059,422 - 7,119,244 - Region in detail - Anopheles gambiae - VectorBase [https://vectorbase.org/vectorbase/app/record/gene/AGAP008222]

33. Chromosome 3R:7,435,306 - 7,485,012 - Anopheles gambiae - VectorBase [https://vectorbase.org/vectorbase/app/record/gene/AGAP008232]

34. Chromosome 3R: 8,176,778 - 8,183,084 - Region in detail - Anopheles gambiae - VectorBase [https://vectorbase.org/vectorbase/app/record/gene/AGAP008273]

35. Bandelt HJ, Forster P, Röhl A: Median-joining networks for inferring intraspecific phylogenies. Mol Biol Evol 1999, 16 (1)37–48.

36. Sabeti PC, Reich DE, Higgins JM, Levine HZP, Richter DJ, Schaffner SF, Gabriel SB, Platko JV, Patterson NJ, McDonald GJ et al: Detecting recent positive selection in the human genome from haplotype structure. Nature 2002, 419 (6909)832–837.

37. Müller P, Warr E, Stevenson BJ, Pignatelli PM, Morgan JC, Steven A, Yawson AE, Mitchell SN, Ranson H, Hemingway J et al: Field-caught permethrin-resistant *Anopheles gambiae* overexpress CYP6P3, a P450 that metabolises pyrethroids. PLoS Genet 2008, 4 (11):e1000286.

38. Stica C, Jeffries CL, Irish SR, Barry Y, Camara D, Yansane I, Kristan M, Walker T, Messenger LA: Characterizing the molecular and metabolic mechanisms of insecticide resistance in Anopheles gambiae in Faranah, Guinea. Malar J 2019, 18 (1):244.

39. Ag1000G - AR3 Panoptes genome browser [https://www.malariagen.net/apps/ag1000g/phase1-AR3/index.html?dataset=Ag1000G&workspace=workspace_1&view=f6c6c7c8-23c9-11eb-a4f3-22000a6287ed&state=genomebrowser]

40. Clarkson CS, Miles A, Harding NJ, Weetman D, Kwiatkowski D, Donnelly M, The Anopheles gambiae Genomes C: The genetic architecture of target-site resistance to pyrethroid insecticides in the African malaria vectors Anopheles gambiae and Anopheles coluzzii. 2018.

41. Hemingway J: The role of vector control in stopping the transmission of malaria: threats and opportunities. Philos Trans R Soc Lond B Biol Sci 2014, 369 (1645):20130431.

42. Liu N: Insecticide resistance in mosquitoes: impact, mechanisms, and research directions. Annu Rev Entomol 2015, 60 537–559.

43. Ingham VA, Pignatelli P, Moore JD, Wagstaff S, Ranson H: The transcription factor Maf-S regulates metabolic resistance to insecticides in the malaria vector Anopheles gambiae. BMC Genomics 2017, 18 (1):669.

44. Wilding CS: Regulating resistance: CncC:Maf, antioxidant response elements and the overexpression of detoxification genes in insecticide resistance. Curr Opin Insect Sci 2018, 27 89–96.

45. Coetzee M, Hunt RH, Wilkerson R, Della Torre A, Coulibaly MB, Besansky NJ: Anopheles coluzzii and Anopheles amharicus, new members of the Anopheles gambiae complex. Zootaxa 2013, 3619 246–274.

46. Holt RA, Subramanian GM, Halpern A, Sutton GG, Charlab R, Nusskern DR, Wincker P, Clark AG, Ribeiro JMC, Wides R et al: The genome sequence of the malaria mosquito Anopheles gambiae. Science 2002, 298 (5591)129–149.

47. Cingolani P, Platts A, Wang LL, Coon M, Nguyen T, Wang L, Land SJ, Lu X, Ruden DM: A program for annotating and predicting the effects of single nucleotide polymorphisms, SnpEff: SNPs in the genome of Drosophila melanogaster strain w1118; iso-2; iso-3. Fly 2012, 6 (2)80–92.

48. Lewontin RC: The Interaction of Selection and Linkage. I. General Considerations; Heterotic Models. Genetics 1964, 49 (1)49–67.

49. Hunter JD: Matplotlib: A 2D Graphics Environment. Comput Sci Eng 2007, 9 (3)90–95.

50. Cartopy: Using cartopy with matplotlib - cartopy 0.18.0 documentation. In 0.17.0 edn. https://scitools.org.uk/; 2020.

51. Wagah MG: ag1000g-phase2-cyp6m2. In., 9/11/2020 edn. https://github.com/; 2020.

52. Harrington B: Inkscape. In., 1.0.1 edn; 2005.

53. Miles A: scikit-allel - Explore and analyse genetic variation - scikit-allel 1.3.2 documentation. In. https://github.com; 2018.

54. Consortium TAgG: Ag1000G phase 2 AR1 data release. In., 1 edn. MalariaGen Genomic Epidemiology Network; 2017.

