## Supplementary figures and images for "Genetic variation at the *Cyp6m2* putative insecticide resistance locus in *Anopheles gambiae* and *Anopheles coluzzii*"

### Supplemental Data 1

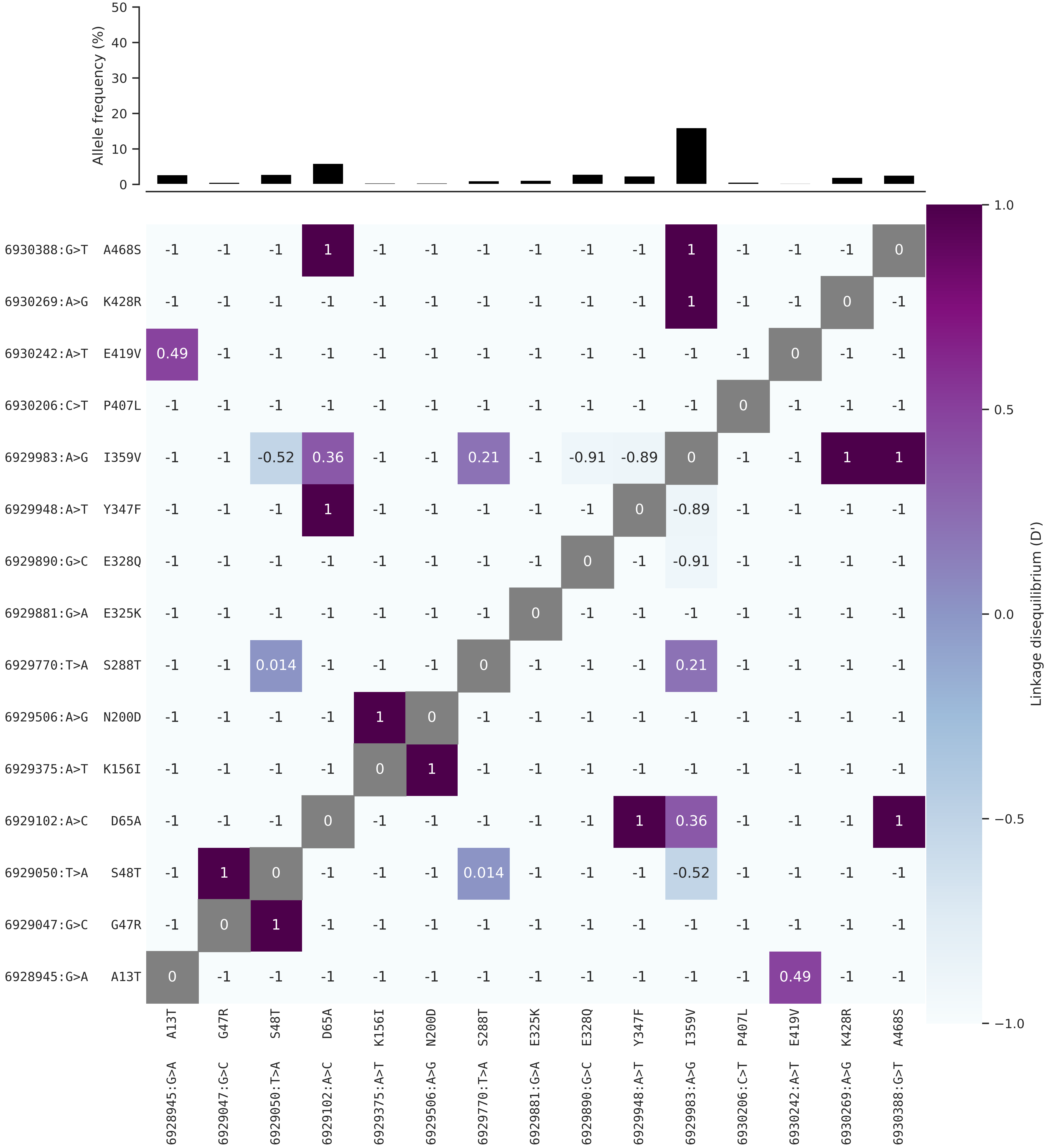

### Supplementary Figure 2. Hierarchical clustering and missense mutations for ODR2.

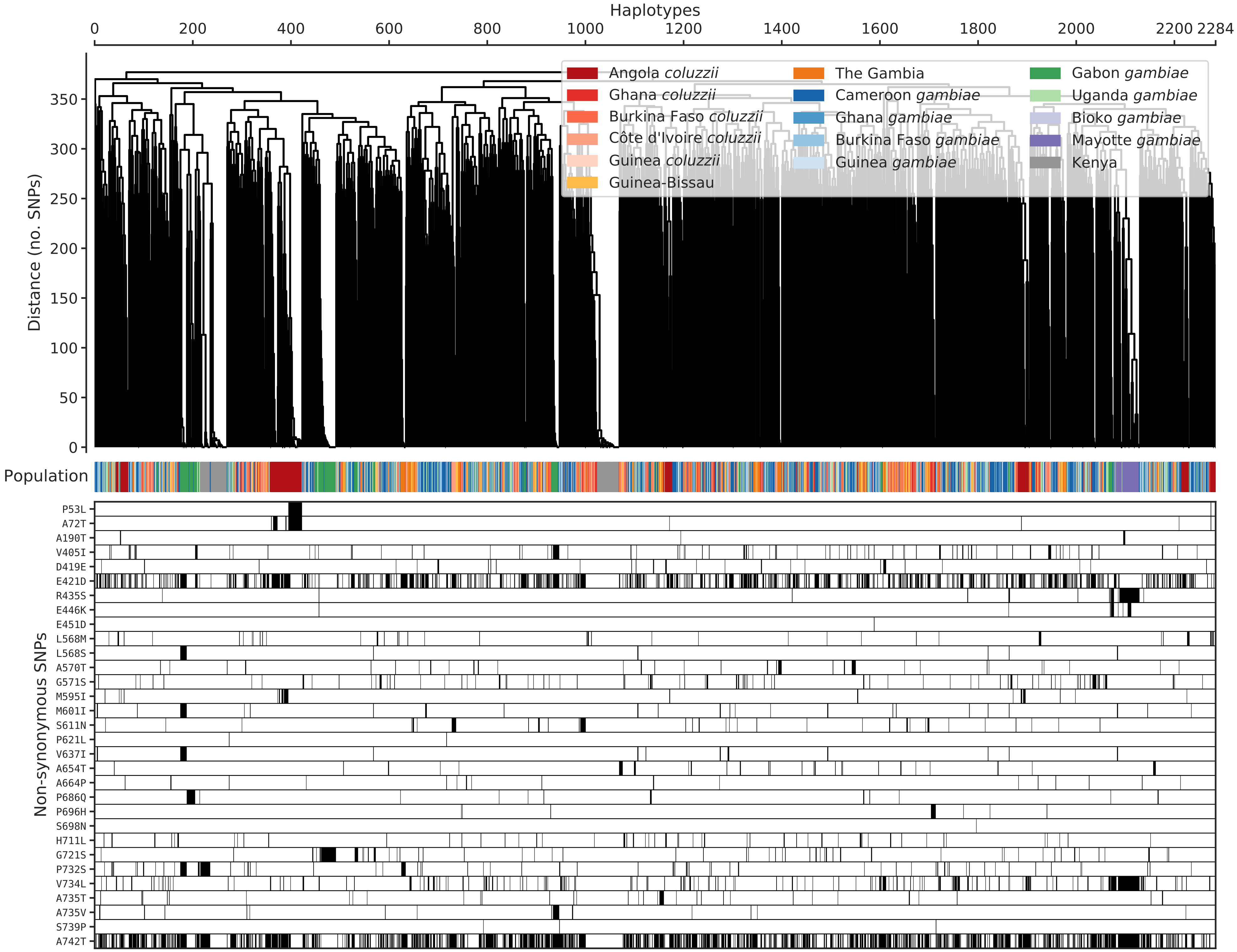

### Supplementary Figure 3. Hierarchical clustering and missense mutations for HAM

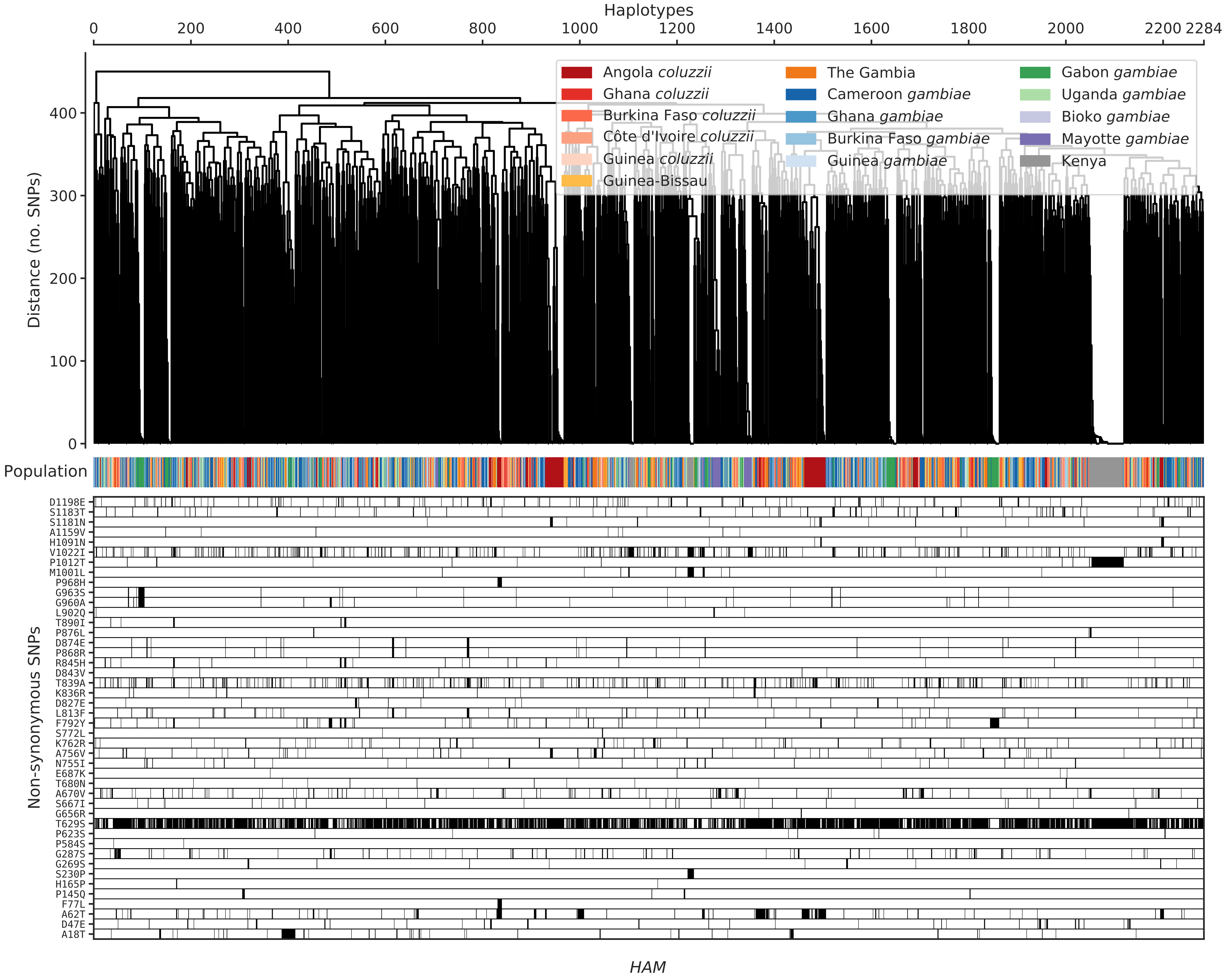

### Supplementary Figure 4. Hierarchical clustering and missense mutations for SH2.

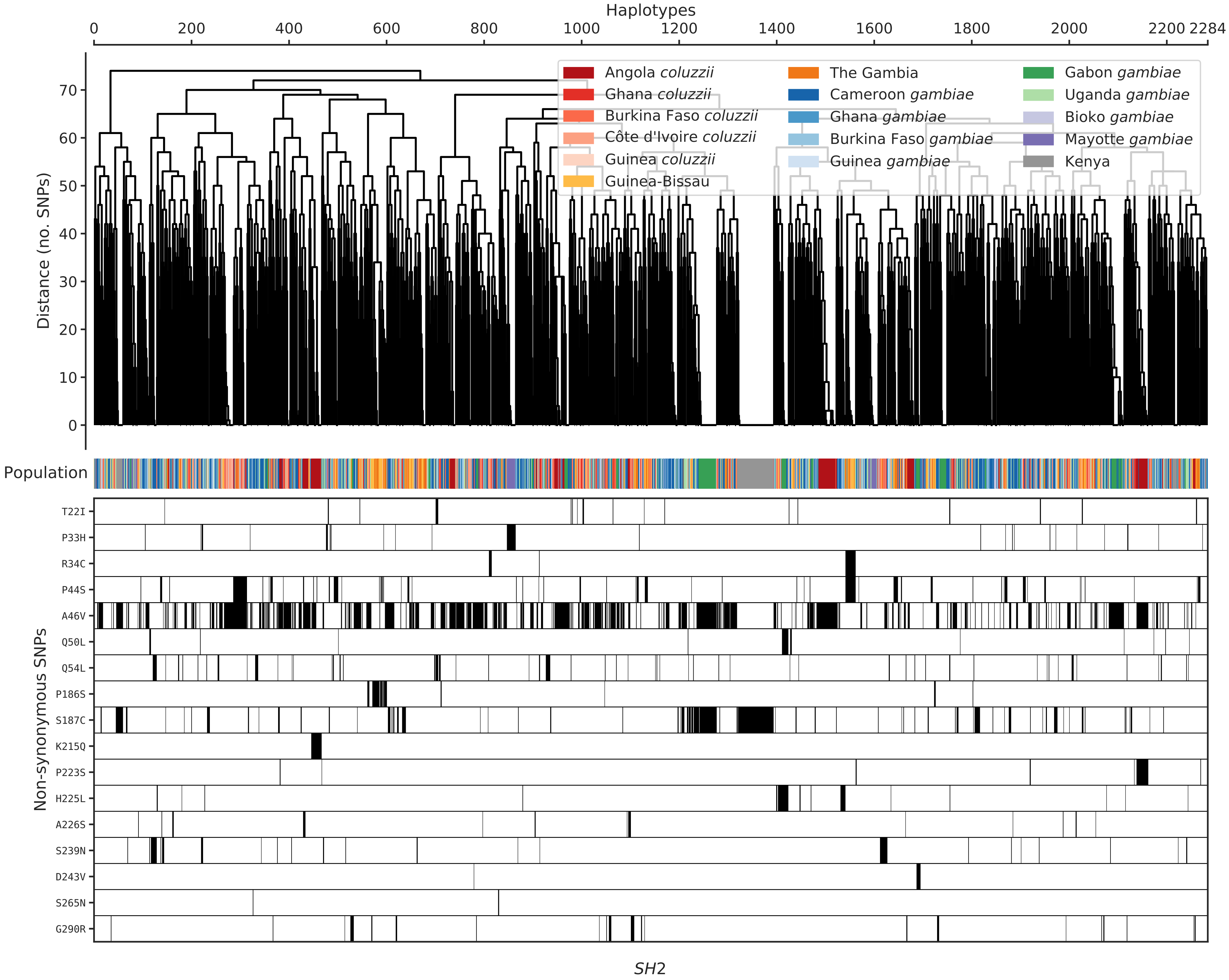

### Supplementary Figure 5. Hierarchical clustering and missense mutations for Cyp6m sub cluster.

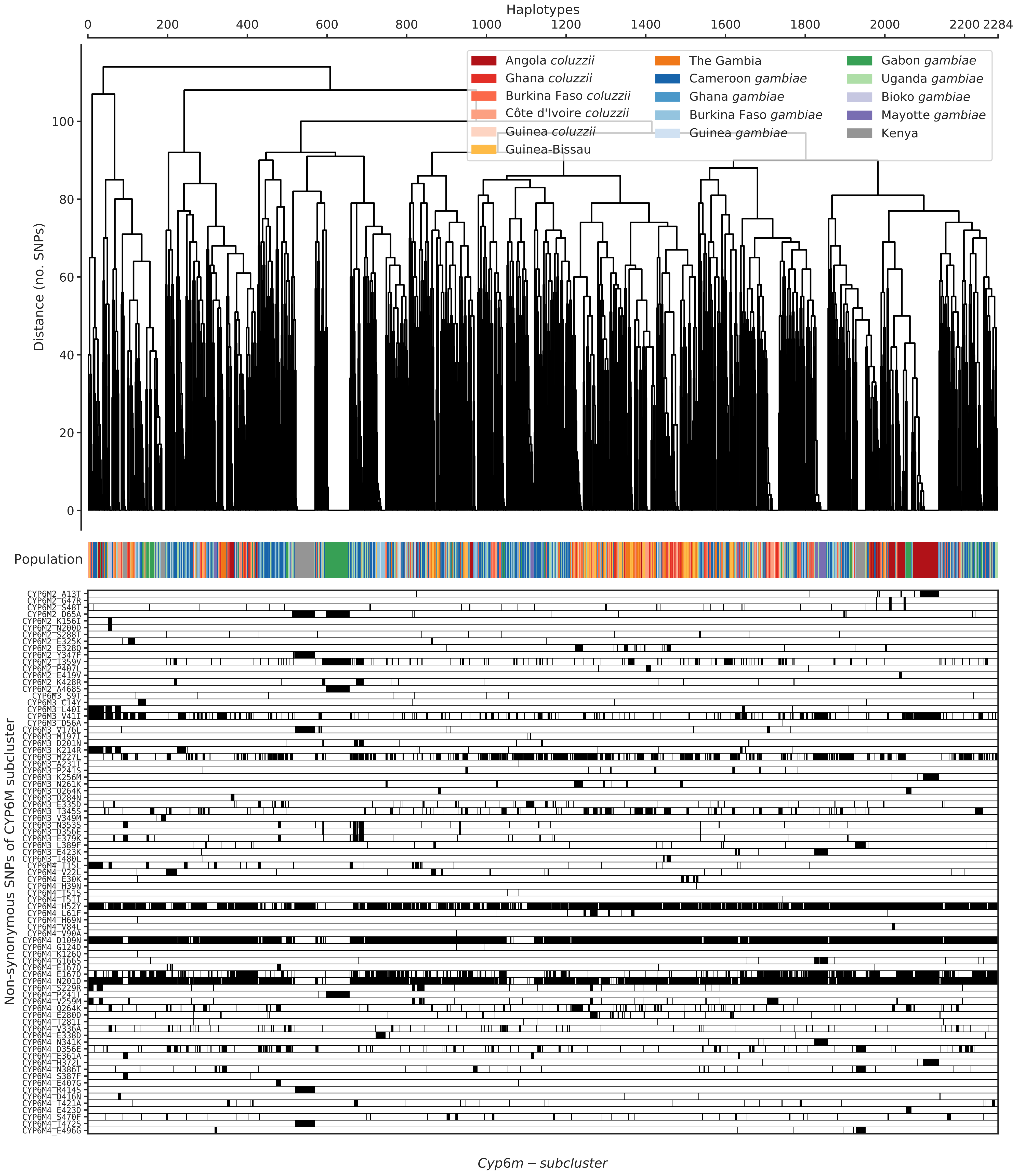

### Supplementary Figure 6. Hierarchical clustering and missense mutations for Cyp6 supercluster.

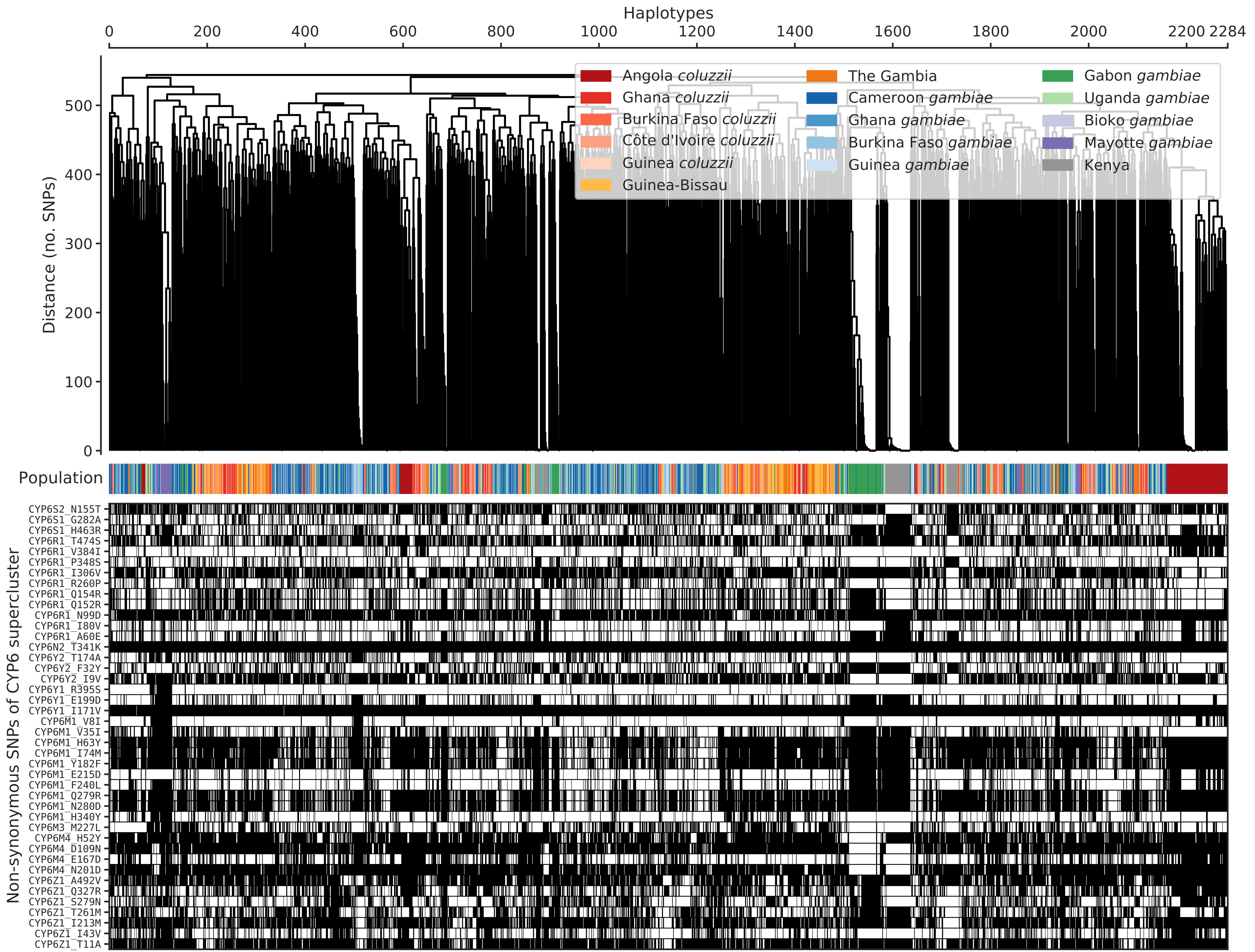


*Cyp6* - supercluster

### Supplementary Figure 7. Extended haplotype homozygosity across all populations.

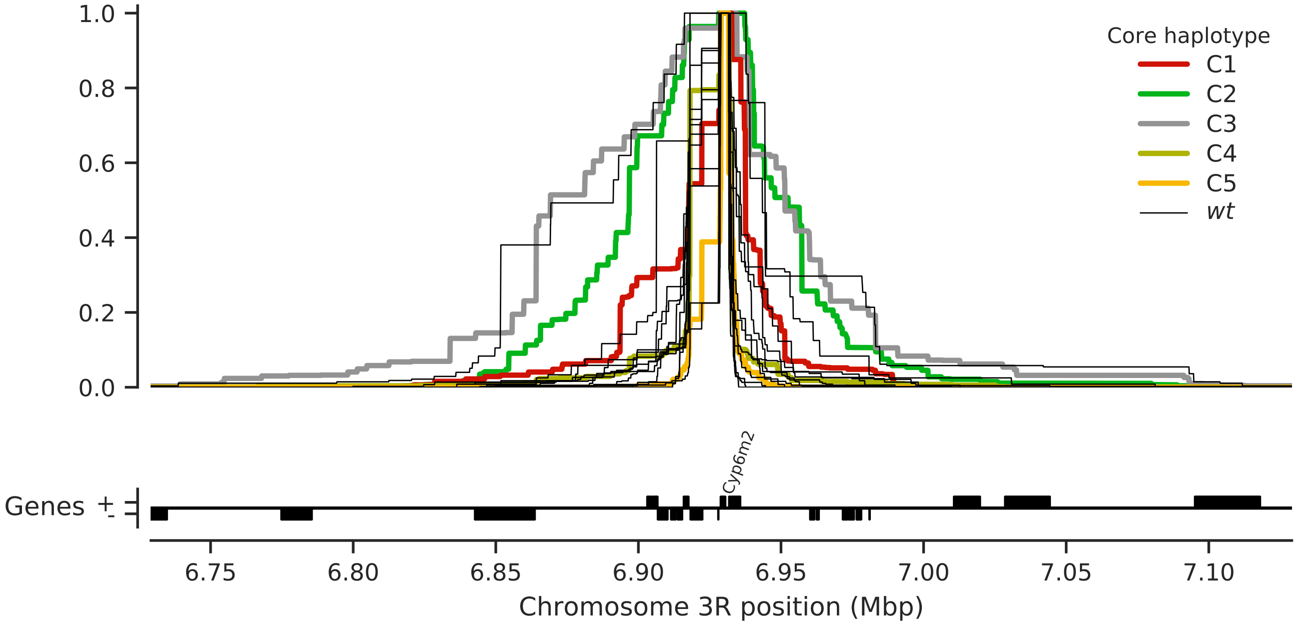
